# Regulatory Role of PKR in Systemic Inflammation-Triggered Neuroinflammation and its Modulation of Glucose Metabolism and Cognitive Functions in Cholinergic Neurons

**DOI:** 10.1101/2023.10.10.561630

**Authors:** Wai-Yin Cheng, Xin-Zin Lee, Michael Siu-Lun Lai, Yuen-Shan Ho, Raymond Chuen-Chung Chang

## Abstract

Systemic inflammation may promote neuroinflammation and neurodegeneration. The double-stranded RNA-dependent protein kinase (PKR) is a key signaling molecule that regulates immune responses. This study aims to examine the role of PKR in regulating systemic inflammation-induced neuroinflammation and cognitive dysfunctions using a laparotomy mouse model. In the first part, wild-type C57BL/6J and C57BL/6-Tg(CD68-EGFP)1Drg/J mice were assigned to undergo either laparotomy with sevoflurane anesthesia or sevoflurane alone to examine effects of systemic inflammation on neuroinflammation and cognition. In the second part, PKR^-/-^ mice were used to study the role of PKR in modulating laparotomy-induced systemic inflammation, neuroinflammation, and cognition. For the third part, PKR was inhibited selectively in cholinergic neurons of ChAT-IRES-Cre-eGFP mice via intracerebroventricular injection of rAAV-DIO-PKR-K296R. This examined the effects of inhibiting PKR in cholinergic neurons on glucose metabolism and cognition in the laparotomy model. Our study revealed that genetic deletion of PKR in mice potently attenuated the laparotomy-induced peripheral and neural inflammation and cognitive deficits. Furthermore, inhibiting PKR in the cholinergic neurons rescued the laparotomy-induced brain glucose hypometabolism and cognitive impairment. Our results demonstrated the critical role of PKR in regulating neuroinflammation and cognitive dysfunctions in a peripheral inflammation model. PKR could be a pharmacological target for treating systemic inflammation-induced neuroinflammation and cognitive dysfunctions.

## Introduction

Systemic inflammation may promote neuroinflammation and neurodegeneration [1–3]. Neuroinflammation contributes to neurodegenerative disorders, including Alzheimer’s disease (AD) [4], Parkinson’s disease (PD) [5], and amyotrophic lateral sclerosis [6]. During systemic inflammation, the infiltration of circulating cytokines and macrophages into the brain can activate microglia [7, 8]. Activated microglia release pro-inflammatory cytokines, leading to neuronal injury and death [9]. These inflammatory mediators can further facilitate the recruitment of leukocyte into the brain, perpetuating neuroinflammation [10].

This study utilizes a nonbacterial endotoxin mouse model of laparotomy to address the systemic inflammation triggered by surgery. This laparotomy model is suitable for investigating perioperative neurocognitive disorder (PND) [11]. As a surgical model, laparotomy [12] outperforms other experimental models including tibial fracture surgery [13–15] and injection of bacteria [16] or lipopolysaccharide (LPS) [17] for studying peripheral inflammation because it allows for a longer period of observation up to 2 weeks [16, 17]. Therefore, the laparotomy model is appropriate for the investigation of whether and how neuroinflammation leads to cognitive dysfunctions upon systemic inflammation.

Double-stranded RNA-dependent protein kinase (PKR) is encoded by the *EIF2AK2* gene, regulates mRNA translation, transcription, proliferation, and apoptosis [18]. Beyond its established antiviral functions, PKR act as a critical signaling molecule modulating immune responses. It regulates macrophage activation [19], different inflammatory pathways, and inflammasome formation [20]. PKR can be activated by cytokines, viral double-stranded RNA (dsRNA), bacterial LPS, and growth factors [20]. PKR activation can regulate activation of various critical inflammatory kinases, including mitogen-activated protein kinases (MAPKs) [21] and inhibitor of nuclear factor-κB (IκB) kinase [22]. Furthermore, PKR also activates and regulates the release of inflammasome-dependent cytokines [23, 24]. Through these myriad functions, PKR critically regulates inflammatory signaling cascades and responses. In addition to regulating immune responses, PKR plays an essential role in cognition and age-related neurodegeneration. Our laboratory was among the first to reveal that PKR activation and eukaryotic initiation factor-2alpha (eIF2α) phosphorylation are implicated in neuronal apoptosis [25]. The overexpression of wild-type PKR in human neuroblastoma cells could enhance the apoptosis induced by β-amyloid peptides; while overexpression of dominant-negative PKR could attenuate the induced apoptosis. Primary cultured neurons from PKR knockout mice showed greater tolerance to β-amyloid neurotoxicity compared with their wild-type counterpart [25]. In addition, knockout of PKR could result in the enhanced late-phase long-term potentiation (LTP) in hippocampal sections, increased network excitability, as well as improved cognitive functions with the enhancement of learning and memory in mice [26]. All these studies suggested the role of PKR in modulating cognitive functions and neurodegeneration.

Considering the dual functions of PKR in modulating immune responses and cognition, we hypothesize that PKR regulates the transduction of systemic inflammation into neuroinflammation, and PKR in cholinergic neurons can modulate cognitive functions. This project aims at investigating whether PKR can be a pharmacological target for preventing systemic inflammation-triggered neuroinflammation and cognitive deficits.

## Methods

### Animal husbandry

Male wild-type C57BL/6J, C57BL/6-Tg(CD68-EGFP)1Drg/J (referred to herein as CD68-eGFP mice), PKR^-/-^, and ChAT-IRES-Cre-eGFP mice (3 months) were obtained from the Centre for Comparative Medicine Research of the University of Hong Kong. This center is fully accredited by the Association for Assessment and Accreditation of Laboratory Animal Care. The mice were housed in cages under controlled temperature (20°C – 22°C), humidity (50% – 60%), with a 12 hour light/dark cycle and ad libitum access to food and water. Mice were allowed to acclimate for 1 week prior to experiment.

### Ethics statement

All experiments were approved by the Department of Health of the Hong Kong SAR government and the Committee on the Use of Live Animals in Teaching and Research of the University of Hong Kong (#4911-18 and #5819-21).

### Experimental protocols

Laparotomy was used as a model to examine neuroimmune responses induced by systemic inflammation. The experimental workflow is summarized in Fig. S1. In part 1, male wild-type C57BL/6J and CD68-eGFP mice were randomized into two groups: laparotomy with sevoflurane anesthesia and control group with sevoflurane anesthesia alone. In part 2, the PKR^-/-^ mice were exposed to laparotomy with sevoflurane anesthesia or sevoflurane anesthesia only to examine the role of PKR in regulating systemic inflammation-triggered neuroinflammation and cognitive deficits. In part 3, rAAV-DIO-PKR-K296R was intracerebroventricularly injected into the right lateral ventricle of ChAT-IRES-Cre-eGFP mice to suppress PKR activation specifically in cholinergic neurons. Mice were then subjected to laparotomy with sevoflurane or sevoflurane anesthesia alone. This allowed examination of the regulatory effects of blocking PKR in cholinergic neurons on glucose metabolism and cognitive function in the laparotomy model.

### Laparotomy

For anesthetization, the animal was placed in a chamber with 5% sevoflurane (Sevorane™, Abbott, Switzerland) and transferred to a nose mask with anesthesia maintained at 3% sevoflurane with a gas flow of 800 ml/min using an inhalation anesthesia machine (Harvard, US). The laparotomy procedure followed previous descriptions with modifications [12]. Hair was removed from the surgical site, and the skin was disinfected. A 2 cm longitudinal midline cut was made at 0.5 cm below the lower right rib. A small intestinal segment of approximately 10 cm length was temporarily exteriorized and subjected to 2 minutes of gentle manipulation before being replaced in the abdomen. Abdominal muscle and skin suture was conducted using 5-0 vicryl and nylon suture respectively (Ethicon, USA).

### Stereotaxic injection

Mice were anesthetized by intraperitoneal injection of a ketamine/xylazine mixture (80 mg/kg and 8 mg/kg, respectively). Prior to surgery, hair was removed from the surgical site, and the skin was disinfected. The head was then fixed onto a stereotaxic frame (Narishige, Japan). A 1.5 cm straight midline incision was made to expose the skull. Either rAAV-DIO-PKR-K296R (packaged in AAV serotype 6) (2 µl; 1×10^13^ vg/mL) or 0.9% saline was injected intracerebroventricularly into right lateral ventricle (antero-posterior: −0.6 mm, medial-lateral: −1.3 mm, dorsal-ventral: −3.1 mm) at a flow rate of 0.5 µL/min using a Hamilton syringe, 33-gauge needle (7803-05, Hamilton, USA). Following the injection, the needle was retained in situ for 5 min allowing the solution to diffuse and avoiding solution backflow. The incision site was sealed with a nonabsorbable nylon suture (5-0, PS-2; Ethicon, USA).

### Open field test

Open field test (OFT) is adopted to assess locomotor activity, exploratory behavior and anxiety in mice based on their innate propensity to avoid open spaces [27]. Mice were placed in a 40 cm square arena and allowed to freely explore for 10 min. Total distance traveled and time spent exploring central area were quantified as indices of locomotor activity and anxiety.

### Novel object recognition test

Novel object recognition test (NOR) assesses recognition memory in mice based on their innate preference for novelty [28]. In the training session, mice were first allowed to explore an open field arena with two identical objects (A + A) for 10 min. After 24 hours, mice were allowed to explore the familiar arena with a novel object replacing one of the identical objects (A + B). Basically, driven by curiosity, the mice would spend more time exploring the novel object rather than the familiar one. The discrimination index, calculated as a ratio of time exploring the novel verus familiar object, can be regarded as an index of recognition memory.

### Spontaneous alternation Y-maze test

Spontaneous alternation Y-maze (SYM) test assesses short-term spatial working memory in rodents. Mice are allowed to explore the maze for 5 min [29]. A spontaneous alternation involves three successive entries into a new arm rather than returning to an earlier one. Driven by curiosity, the mice will tend to explore the new arm rather than the previously visited arm. The percentage of alternation was determined by dividing the number of spontaneous alternations by the total arm entries, then multiplying by 100. The percentage of alternation can be the indicator of short-term spatial working memory.

### Puzzle box test

A puzzle box test was a five-day protocol modified by our laboratory [28] based on the previous published method [30]. It is designed to assess rodent’s native problem-solving ability, short-term and long-term memory. The puzzle box was consisted of two main sections: an open-field area (600 × 280 mm) and an adjacent sheltered goal-box area (600 × 280 mm). Mice could enter the sheltered goal-box arena through a small underpass between these two compartments. Mice were required to complete escape tasks of increasing difficulty within a fixed period of time (Table S1). The first four days involved three trials per day, while the final day comprised a single trial. Each obstruction includes three trials; the initial two taking place on the same day, with the third trial conducted on the following day. Three trials were performed: the first evaluated problem-solving skills; the second assessed short-term memory, and the third investigated long-term memory.

### Arrangement of behavioral tests

A series of cognitive behavioral tests was conducted based on the following schedule. For minimized fatigue and interference, behavioral tests were divided across the study. A group of mice underwent OFT on postoperative day (POD) 10, followed by NOR on POD 11–12. A second group of mice underwent OFT on POD 10, followed by SYM on POD 11. The third cohort completed OFT on POD 9 and the puzzle box test on POD 10–14.

### RNA extraction and quantitative real-time RT-PCR

Mice were sacrificed by carbon dioxide asphyxiation and decapitation, in line with the protocols recommended by American Veterinary Medical Association and then followed by transcardial perfusion with ice-cold saline. Frontal cortex, hippocampus, and liver were harvested, snap-frozen, and kept at −80°C. Total RNA was isolated from samples using the RNAiso Plus (TAKARA, Japan). Detailed protocols were described in Supplementary Methods. Quantitative real-time RT-PCR was performed with TB Green® Premix Ex Taq™ II (TAKARA, Japan) using a Bio-Rad CFX 96 real-time PCR detection system (Bio-Rad, USA) in triplicate. Primer sequences utilized for amplification are provided in Table S2. Relative expression was determined using the comparative CT method, normalizing the target gene mRNA to GAPDH.

### Immunohistochemical staining and confocal microscopy

The processes for cryosection preparation described in the Supplementary Methods were similar to those described previously for human brain sections [31]. Prior to confocal imaging, brain slices were pretreated with 10% goat serum blocking solution, followed by overnight incubation at 4°C with primary antibodies listed in Table S3, followed by 1h secondary antibody staining (Alexa Fluor, 1:400, Invitrogen) at room temperature and co-staining with 4[, 6-diamidino-2-phenylindole (DAPI). Confocal images were acquired using a LSM800 confocal microscope (Carl Zeiss, Germany) with 10× or 20× objective or 40× oil immersion objective using Zen Blue software. The frame size was set to 1024 × 1024 pixels, the bit depths to 16-bit, averaging to 2×, and pinhole setting at 1 airy unit. Z-stack images were acquired, followed by orthogonal projections from Z-stack images using Zen Blue software. Image analysis was performed using ImageJ software (National Institute of Health, USA).

### Flow cytometry

The frontal cortex and hippocampus of CD68-eGFP mice were collected, processed through homogenization and filtration to obtain single-cell suspensions, which were then stained with DAPI. Detailed protocol was described in the Supplementary Methods. Cellular fluorescence was measured using an Agilent NovoCyte Quanteon analyzer (Agilent, USA).

### Targeted metabolomics by GC-MS/MS

Samples were obtained from the collected frontal cortex and prepared using the protocol described in the Supplementary Methods. Targeted metabolomics was performed on central carbon metabolism using gas chromatography-tandem mass spectrometry (GC-MS/MS) on Agilent 7890B GC-Agilent 7010 Triple Quadrupole Mass Spectrometer system (Agilent, USA). The experimental parameters were stated in the Supplementary Methods. Data were normalized by median and log-transformed. Heat map and hierarchical clustering were conducted based on Euclidian distance using Ward’s algorithm. Further principal component analysis (PCA) and partial least squares discriminant analysis (PLS-DA) were further performed using Metaboanalyst 5.0.

### Statistical analysis

Unless otherwise specified, all statistical analyses were performed on Prism v8.0c (GraphPad Software, USA). Data were expressed as means ± SEM. The normality of data were assessed using the D’Agostino-Pearson normality test. Unpaired two-tailed Student’s t-test at *p* < 0.05 was applied for comparing two group to determine the statistical significance of treatment effects. Interactions between mouse strain and treatment were assessed using two-way ANOVA with Bonferroni post hoc test. Statistical significance was set at *p* < 0.05.

## Results

### Laparotomy induced peripheral and neural inflammation in wild-type mice

To investigate the systemic and neural immune responses following the laparotomy challenge, we initially performed laparotomy in the male C57BL/6J mice. Four hours following surgery, IL-1β and monocyte chemoattractant protein (MCP)-1 mRNA expression were enhanced in the liver of the laparotomy group when compared with that in the control counterpart (*p* < 0.05; Fig. 1a). IL-1β expression was also significantly upregulated in the hippocampus following the laparotomy challenge (*p* < 0.05; Fig. 1a). On POD 1, no significant differences in IL-1β, IL-6, MCP-1, and TNF-α mRNA expression in the liver were found between the laparotomy and control groups (Fig. 1b). This finding implied that the induced systemic inflammation is relatively transient. However, laparotomy significantly increased IL-1β and IL-6 expression in the frontal cortex and hippocampus versus controls (*p* < 0.05; Fig. 1b), indicating ongoing, persistent neuroimmune responses.

**Fig. 1.**
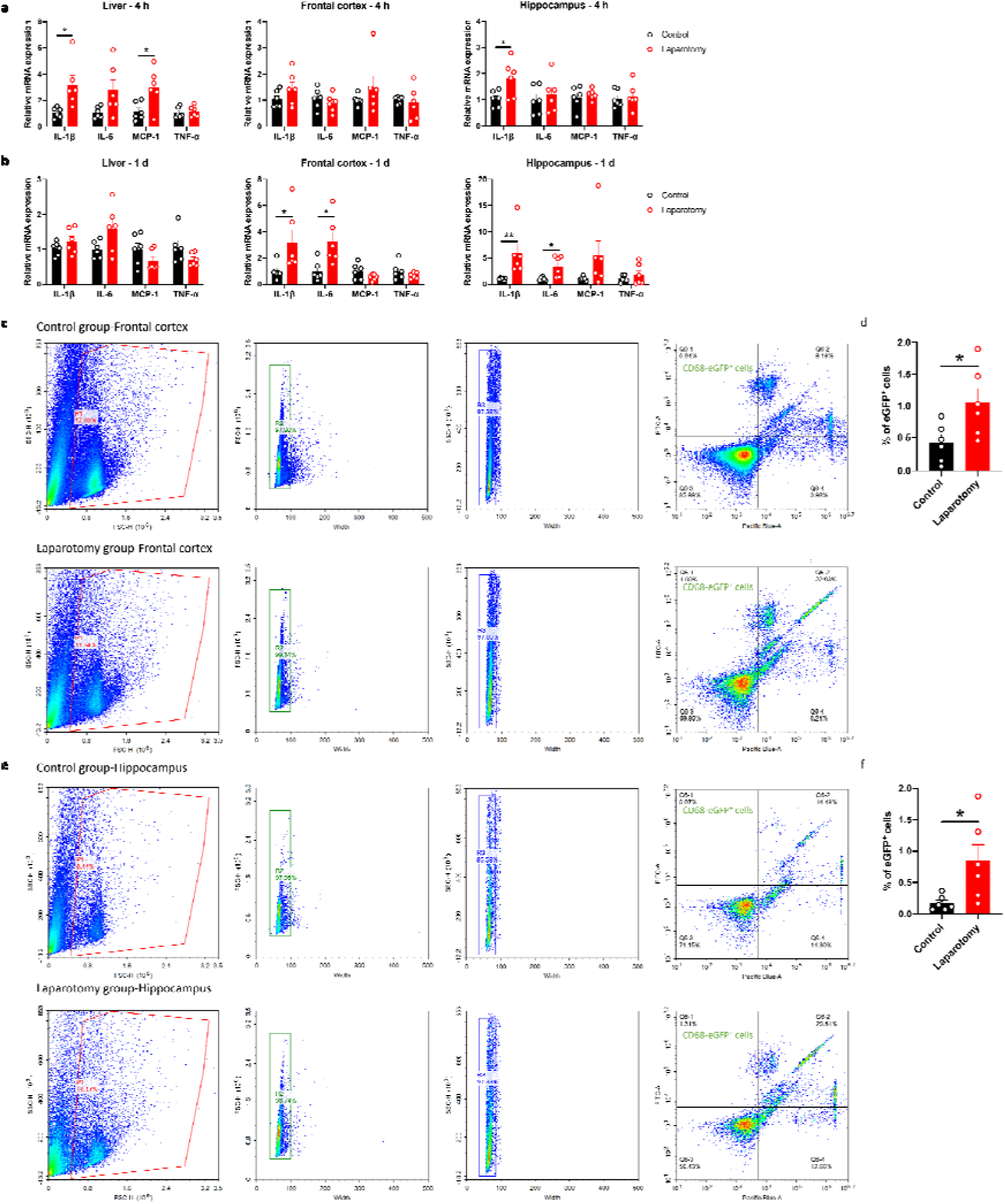
Laparotomy induced systemic inflammation and neuroinflammation. Relative cytokine mRNA expression levels were measured in the liver, frontal cortex, and hippocampus of C57BL/6J mice at (A) 4 h and (B) 1 day after laparotomy. Representative flow cytometry plots illustrating the increase in CD68-eGFP^+^ cells in the (C) frontal cortex and (E) hippocampus of the CD68-eGFP mice on postoperative day 14. Morphological gating strategy refers to dot plot SSC-H versus FSC-H to eliminate debris and aggregates in the (C) frontal cortex and (E) hippocampus of the control and laparotomy groups. Doublets are excluded by plotting FSC-H versus width and SSC-H versus width. Proportion of CD68-eGFP^+^ cells (labeled with eGFP) and cell viability (labeled with DAPI) were determined by flow cytometry with FITC and pacific blue channel respectively. The data generated are plotted in two-dimensional dot plots in which FITC-A versus Pacific-Blue-A. A four-quadrant gate was established: Q1) live, CD68-eGFP^+^ cells (FITC positive/Pacific blue negative); Q2) dead, CD68-eGFP^+^ cells (FITC positive/Pacific blue positive); Q3) live cells without CD68 expression (FITC negative/Pacific blue negative); and Q4, dead cells without CD68 expression (FITC negative/Pacific blue positive). Percentage of CD68-eGFP^+^ cells in the (D) frontal cortex and (F) hippocampus of the control and laparotomy groups. n = 6 per group. Data are expressed as mean ± S.E.M. Differences were assessed by unpaired two-tailed Student’s t-test as follows: \**p* < 0.05 and \*\**p* < 0.01 compared with the control group.

To further study the effects of laparotomy on neuroinflammation, we utilized transgenic CD68-eGFP mice for visualizing microglia/macrophage dynamics. The CD68-eGFP mice expressed high levels of enhanced green fluorescent protein (eGFP) in macrophages and activated microglia, which aided in identifying activated microglia and macrophages. Brain sections were immunostained with antibodies against eGFP, TMEM119 (transmembrane protein 119: a microglia-specific marker), and CD105 (endoglin: a vascular endothelial marker). On POD 14, more CD68-EGFP-positive cells were observed in the frontal cortex (*p* < 0.01) and hippocampus (including cornu ammonis (CA)1 and CA3 (*p* < 0.05), as well as dentate gyrus (DG) (*p* < 0.01)) of the laparotomy mice compared with those of the control counterpart (Fig. S2, S3). Furthermore, an increased number of vessel-associated macrophages (CD68-eGFP^+^, TMEM119^−^ cells associated with the blood vessels) in the frontal cortex was detected following laparotomy (*p* < 0.05; Fig. S2a, b). This result suggested the activation of microglia/potential infiltration of macrophages following laparotomy. Flow cytometry also demonstrated elevated numbers of CD68-eGFP-positive cells in both frontal cortex and hippocampus on POD 14 when compared to controls (*p* < 0.05; Fig. 1c–f), suggesting laparotomy induced the activation of microglia/potential infiltration of macrophages in the brain.

Brain sections from the CD68-eGFP mice were immunostained with a microglial activation marker, ionized calcium-binding adaptor molecule 1 (Iba1), to further confirm whether laparotomy induces activation of microglia. More Iba1^+^ cells were found in both frontal cortex and hippocampus of the laparotomy group when compared to controls (Frontal cortex: *p* < 0.05; CA1 and CA3: *p* < 0.001; DG: *p* < 0.01; Fig. 2). Increased fluorescence intensity was also observed in the hippocampus following laparotomy (*p* < 0.05; Fig. 2). Microglial morphology, an important indicator of neuroinflammation, was further quantified by skeleton analysis [32]. Reduced total endpoints or/and shorter process length per microglial cell were observed in the frontal cortex and the CA1 and DG of the hippocampus of the laparotomy group compared with those of the control counterpart (p < 0.05; Fig. 2b, d and h). This finding revealed that microglia were deramified following laparotomy.

**Fig. 2.**
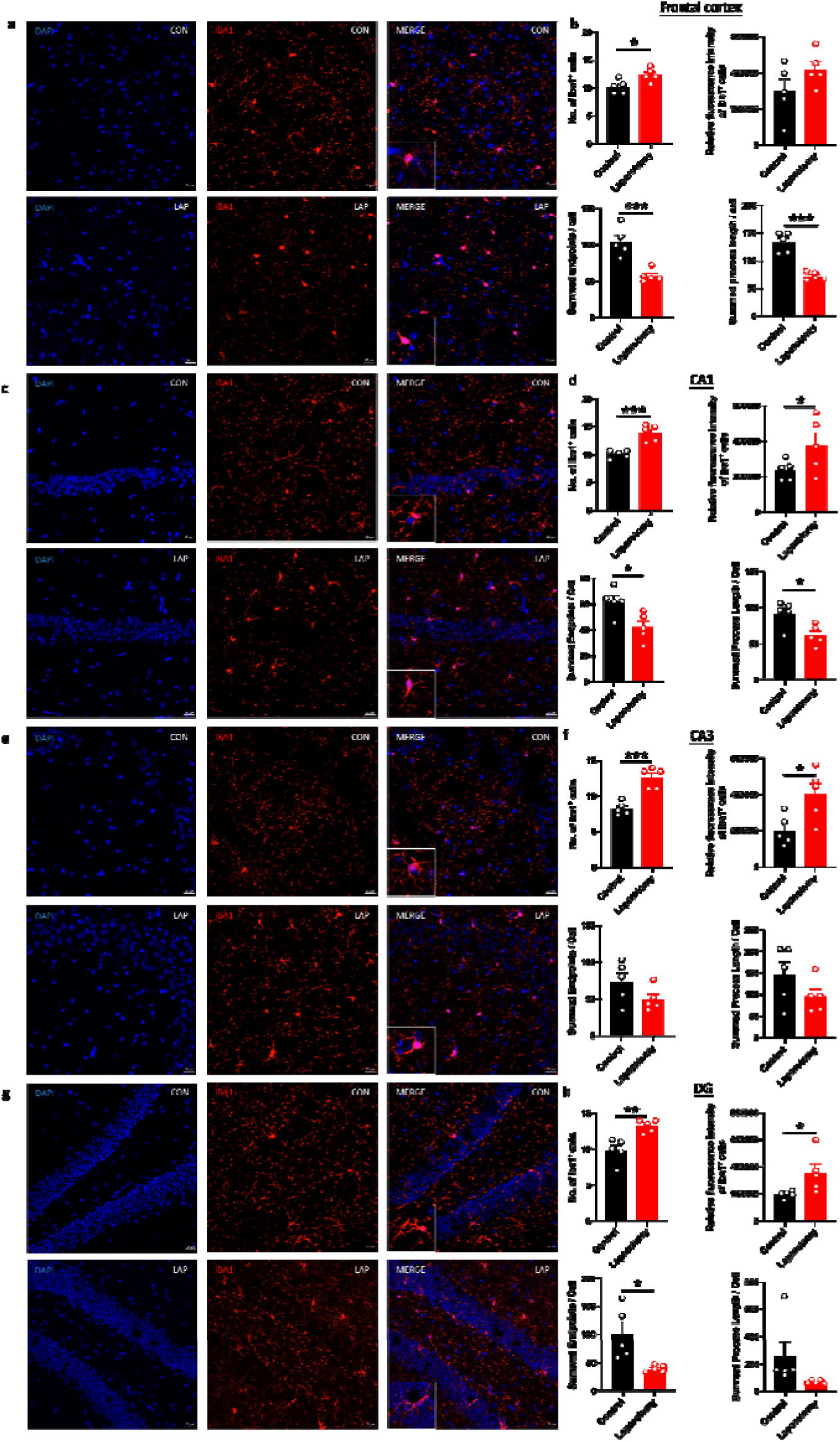
Laparotomy induced activation of microglia in the frontal cortex and hippocampus after laparotomy. Representative confocal images of the immunohistochemical staining of DAPI, and Iba1 in the (A) frontal cortex, (C) cornu ammonis (CA) 1, (E) CA3, and (G) dentate gyrus (DG) of the hippocampus sections from CD68-eGFP mice of the control and laparotomy groups on postoperative day 14. Number of Iba1 positive cells, relative fluorescence intensity of Iba1 positive cells, total number of endpoints per Iba1 positive cell, and summed process length per Iba1 positive cell in the (B) frontal cortex, (D) CA 1, (F) CA3, and (H) DG of the hippocampus sections were quantified. n = 5 per group. Data are expressed as mean ± S.E.M. Differences were assessed by unpaired two-tailed Student’s t-test denoted as follows: \**p* < 0.05, \*\**p* < 0.01 and \*\*\**p* < 0.001 compared with the control group.

### Changes of body weight and cognitive performance in wild-type mice following laparotomy

Body weights were recorded throughout the experiment. Wild-type laparotomy mice exhibited notable weight loss in the first 6 days following surgery (*p* < 0.05; Fig. 3a) but no remarkable decreases thereafter.

**Fig. 3.**
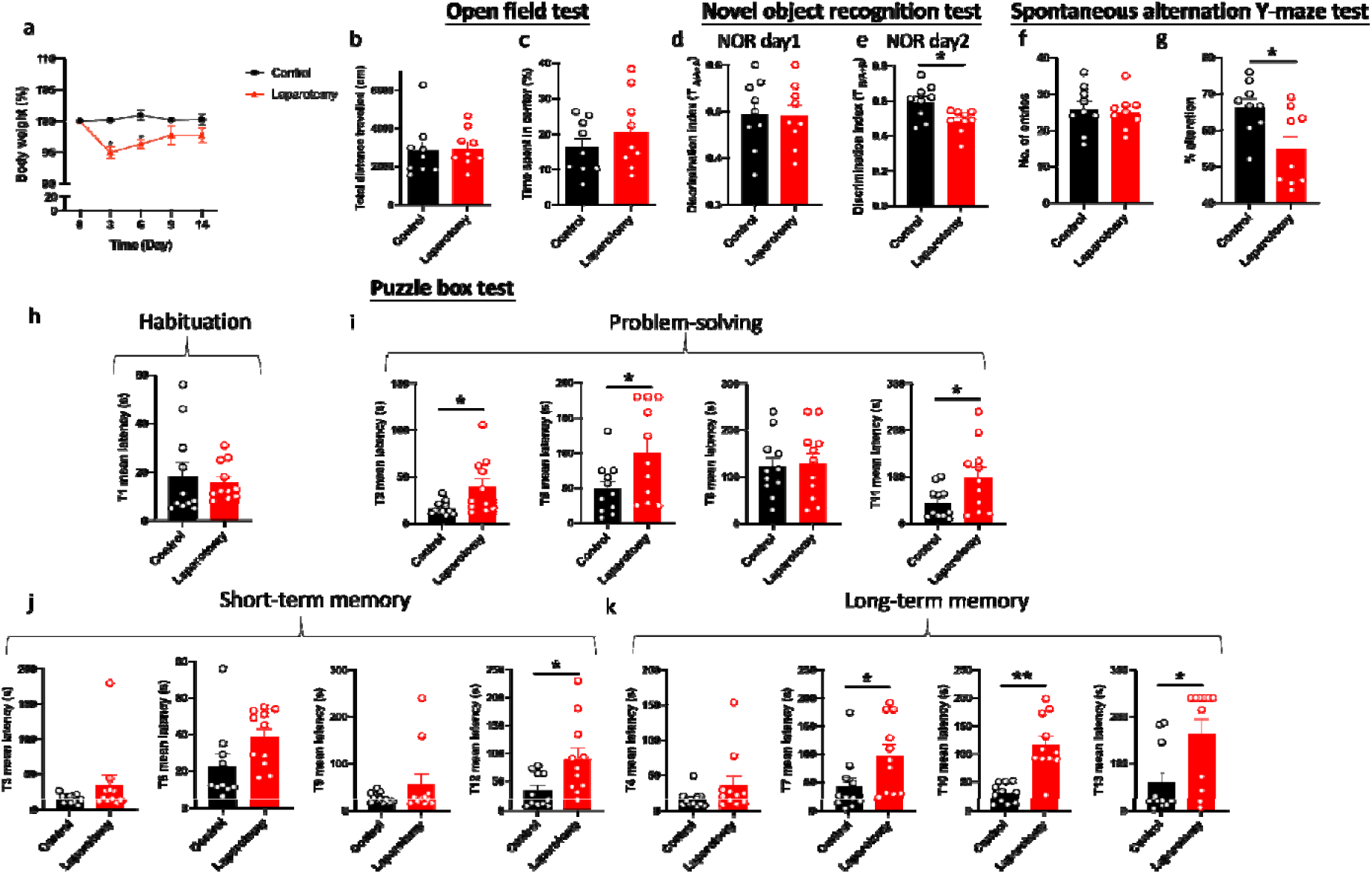
Cognitive impairment of C57BL/6J mice following laparotomy. (A) Percentage of the body weight change from baseline of the laparotomy and control groups throughout the postoperative period. (B) Total distance travelled and (C) central duration of mice in the open-field test. (D) Discrimination index, which is the ratio of exploration time of one object to two objects, was measured on the novel object recognition test day 1 (with two identical objects: A+A) and (E) day 2 (with a novel object replaced the familiar one: A+B). (F) Total number of arm entries and (G) percentage of alternation were measured in the spontaneous alternation Y-maze test. In the puzzle box test, the latency time for the mice entering the goal zone in different tasks was measured. All the tasks include (H) task for habituation (T1), (I) tasks for testing problem-solving skills (T2, T5, T8, and T11), and (J) tasks for evaluating short-term (T3, T6, T9, and T12) and (K) long-term memories (T4, T7, T10, and T13) in the puzzle box test. n = 9–11per group. Data are expressed as mean ± S.E.M. Differences were assessed by unpaired two-tailed Student’s t-test denoted as follows: \**p* < 0.05 and \*\**p* < 0.01 compared with the control group.

A series of behavioral tests was performed to study whether laparotomy affects cognitive functions in the wild-type mice. During the OFT, total distance traveled and central area exploration time were similar between laparotomy and control groups (Fig. 3b, c). This finding implied that laparotomy did not affect the locomotor activity, exploration, or anxiety. NOR then evaluated recognition memory according to the innate preference of mice for novelty [28]. On the first day of NOR, both groups of mice spent comparable time exploring the identical objects (Fig. 3d). On day 2 of NOR, impaired recognition memory with lower discrimination index was observed in the laparotomy group versus the controls (*p* < 0.05; Fig. 3e). As indicated by the reduced percentage of alternation, impaired spatial working memory was detected in the laparotomy group when compared to the control (*p* < 0.05; Fig. 3g) during SYM with similar total number of entries (Fig. 3f). Puzzle box test was adopted to examine the rodent’s native problem-solving ability during trial (T)2, T5, T8, and T11. Additionally, evaluations at the T3, T6, T9, and T12 provided insights into their short-term memory recall, while the T4, T7, T10, and T13 were indicative of their long-term memory function [28]. In the habituation task (T1), both groups spent similar time to complete the escape task (Fig. 3h). Impaired problem-solving abilities requiring a longer period of time to complete the escape task in T2, T5, and T11 were detected in the laparotomy group compared to controls (*p* < 0.05; Fig. 3i). Furthermore, the laparotomy group showed reduced short-term (with increased latency in T12) (*p* < 0.05; Fig. 3j) and long-term (with increased latency in T7, T10, and T13) memories when compared to controls (T7 and T13: *p* < 0.05; T10: *p* < 0.01; Fig. 3k).

### Knockout of PKR in mice potently downregulated the up-regulated pro-inflammatory cytokine expressions induced by laparotomy and ameliorated the microglial activation following laparotomy

Given that PKR regulates immune responses, we assessed whether genetic deletion of PKR could abrogate laparotomy-induced inflammation. IL-1β expression was significantly elevated in liver and hippocampus of the wild-type mice at four hours post-surgery (*p* < 0.05; Fig. 4a) but not in the PKR^-/-^ mice after the surgery compared with that in the controls (Fig. 4a). Meanwhile, expression of IL-1β in liver and frontal cortex was significantly reduced in the PKR^-/-^ laparotomy group when compared to the wild-type laparotomy group (*p* < 0.05; Fig. 4a). With regard to MCP-1 expression in the liver, a significant increase was observed in the wild-type mice after laparotomy (*p* < 0.05; Fig. 4a) while not in the PKR^-/-^ mice (Fig. 4a). Hepatic TNF-α expression was significantly elevated in PKR^-/-^ mice after laparotomy compared to PKR^-/-^ controls (*p* < 0.05; Fig. 4a); wild-type mice showed no such difference.

**Fig. 4.**
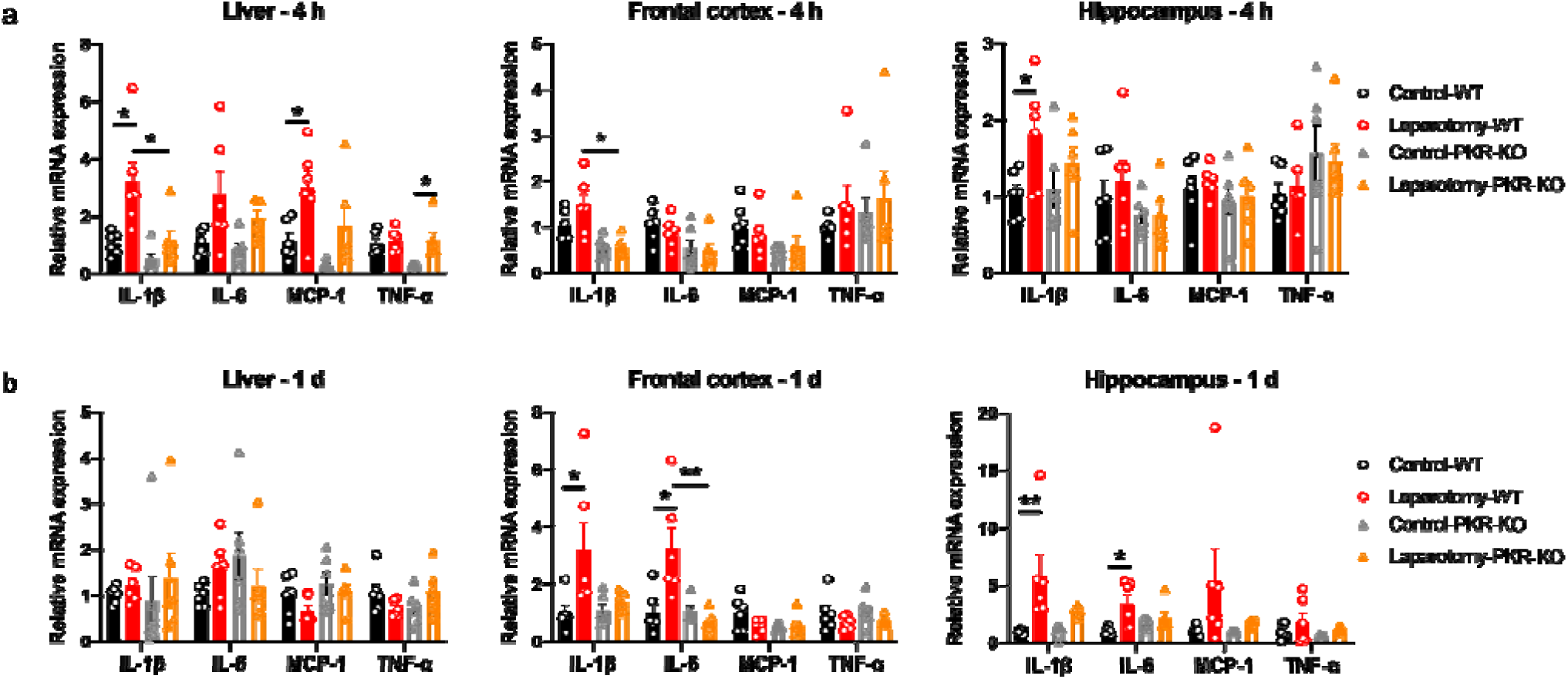
Knockout of PKR in mice ameliorated laparotomy-induced peripheral and neural inflammation after surgery. Relative cytokine mRNA expression levels were measured in the liver, frontal cortex, and hippocampus of the C57BL/6J mice and PKR-KO mice at (A) 4 h and (B) 1 day following laparotomy. n = 6 per group. Data are expressed as mean ± S.E.M. Differences were assessed by two-way ANOVA and Bonferroni’s post hoc test denoted as follows: \**p* < 0.05 and \*\**p* < 0.01, compared with the control group.

On POD 1, the wild-type laparotomy group revealed a substantial increase in IL-1β and IL-6 mRNA expression in frontal cortex and hippocampus, compared to the wild-type controls (*p* < 0.05; Fig. 4b); no such increases were detected in the PKR^-/-^ laparotomy group. Meanwhile, the PKR^-/-^ laparotomy group showed significantly lower IL-6 expression in the frontal cortex versus the wild-type laparotomy group (*p* < 0.01; Fig. 4b). Between the surgical and control groups for both strains of mice, no remarkable differences in IL-1β, IL-6, MCP-1, and TNF-α mRNA expression were detected in the liver (Fig. 4b) and no remarkable alterations in MCP-1 and TNF-α expression were found in frontal cortex and hippocampus (Fig. 4b).

We then further investigated whether knockout of PKR could inhibit activation of microglia in the laparotomy model. No observable difference in the number and fluorescence intensity of Iba1^+^ cells in the frontal cortex (Fig. 5a and b) and hippocampus was found between the PKR^-/-^ laparotomy group and its control counterpart (Fig. 5c–h). Concurrently, laparotomy did not induce morphological changes (total number of endpoints and summed branch lengths) in microglia in the frontal cortex (Fig. 5a and b) and hippocampus (Fig. 5c–h) of PKR^-/-^ mice compared with those in the control. The PKR^-/-^ mice showed ameliorated neuroimmune responses compared to wild-type mice, indicating the critical role of PKR in regulating neuroinflammation following laparotomy.

**Fig. 5.**
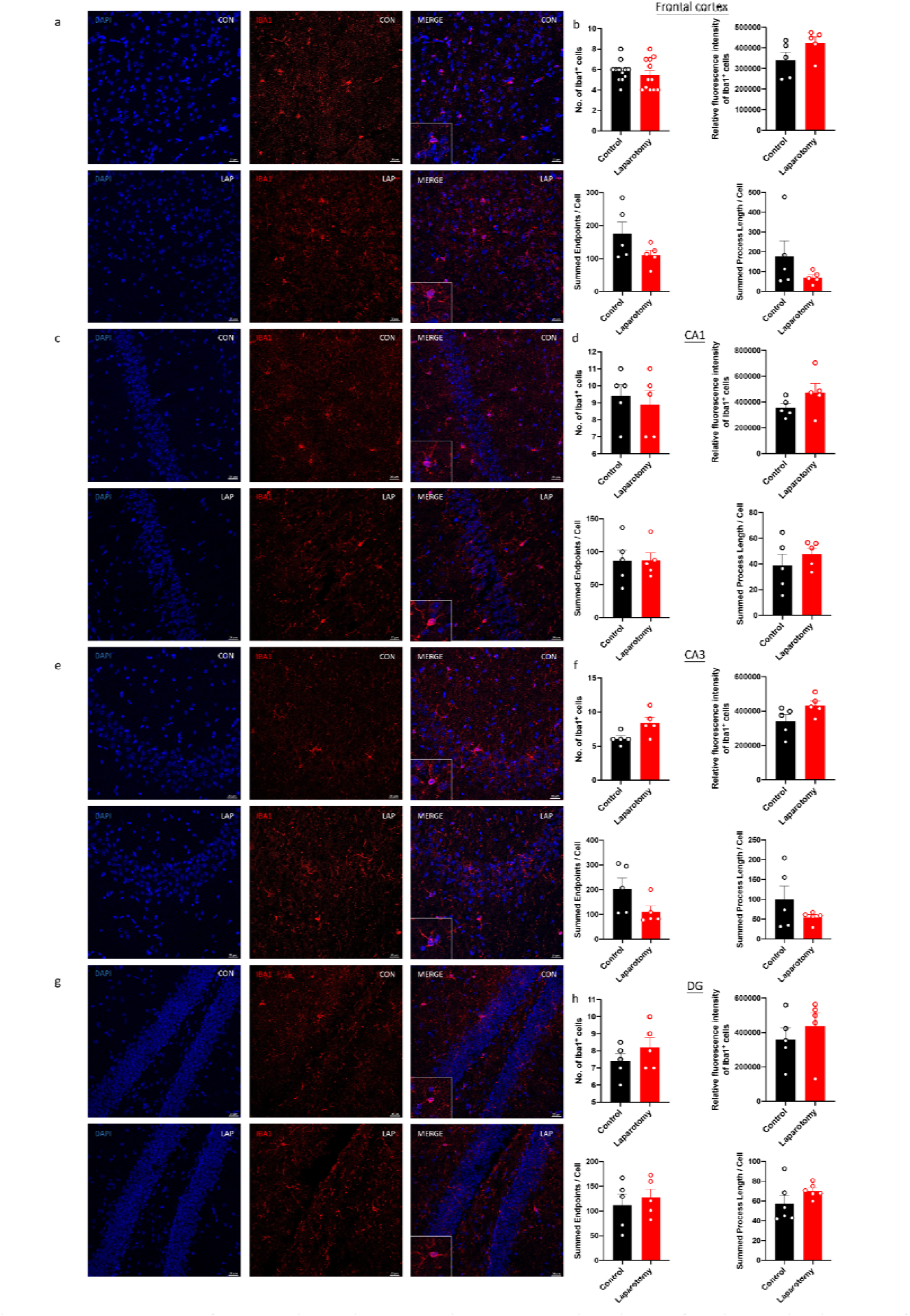
Knockout of PKR in mice ameliorated activation of microglia in the frontal cortex and hippocampus following laparotomy. Representative confocal images of the immunohistochemical staining of DAPI and Iba1 in the (A) frontal cortex, (C) cornu ammonis (CA) 1, (E) CA3, and (G) dentate gyrus (DG) of the hippocampus sections from PKR-KO mice of the control and laparotomy groups on postoperative day 14. Number of Iba1 positive cells, relative fluorescence intensity of Iba1 positive cells, total number of endpoints per Iba1 positive cell, and summed process length per Iba1 positive cell in the (B) frontal cortex, (D) CA1, (F) CA3, and (H) DG of the hippocampus sections were quantified. n = 5 per group. Data are expressed as mean ± S.E.M. Differences were assessed by unpaired two-tailed Student’s t-test.

### Knockout of PKR in mice ameliorated the cognitive deficits induced by laparotomy

Regardless of the mouse strain, laparotomy resulted in a considerable body weight loss within 3 days (*p* < 0.05; Fig. 6a). PKR^-/-^ mice restored weight by day 6, earlier than that of the wild-type mice (on day 9). In the laparotomy group, neither strain of mice lost significant weight after day 9 (Fig. 6a). Meanwhile, laparotomy did not affect the locomotor activity (total distance travelled) and anxiety level (central duration) of the PKR^-/-^ mice (Fig. 6b and c). On NOR day 1, both strains spent comparable time exploring the identical objects (Fig. 6d). On NOR day 2 with a novel object, laparotomy impaired the recognition memory (reduced discrimination index) of the wild-type mice (*p* < 0.05; Fig. 6e) but not that of the PKR^-/-^ mice (Fig. 6e). In the SYM, laparotomy impaired the spatial recognition memory (reduced percentage of alternation) of the wild-type mice (*p* < 0.05; Fig. 6g) but not that of the PKR^-/-^ mice (Fig. 6g). No notable difference in the total entries was detected between the surgery and control groups (Fig. 6f).

**Fig. 6.**
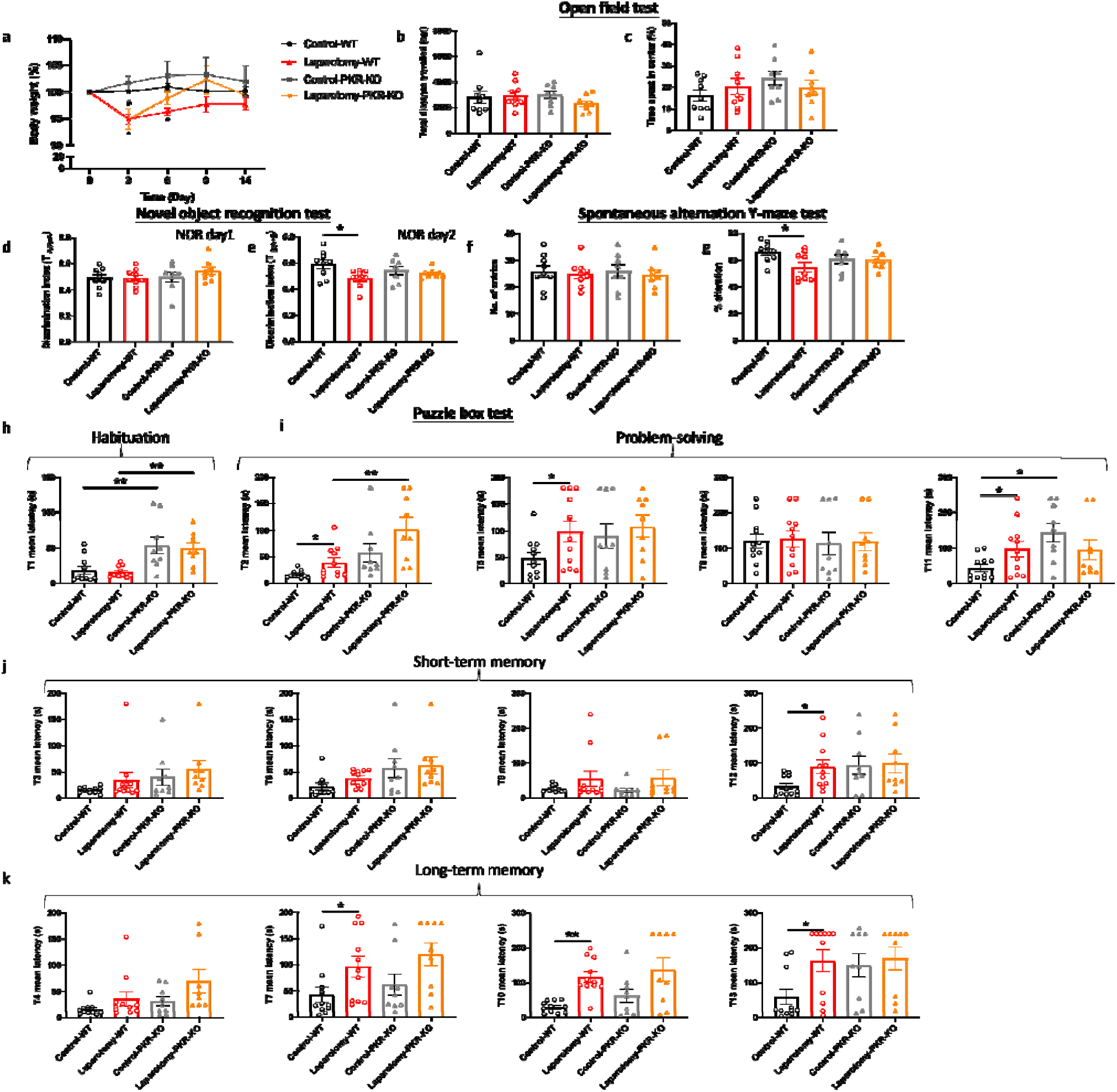
Genetic deletion of PKR in mice alleviated laparotomy-induced cognitive impairment. (A) Percentage of the body weight change from baseline of the C57BL/6J mice and PKR-KO mice throughout the postoperative period. (B) Total distance travelled and (C) central duration of mice in the open-field test. (D) Discrimination index, which is the ratio of exploration time of one object to two objects, was measured on the novel object recognition test day 1 (with two identical objects: A+A) and (E) day 2 (with a novel object replaced the familiar one: A+B). (F) Total number of arm entries and (G) percentage of alternation were measured in the spontaneous alternation Y-maze test. In the puzzle box test, the latency time for the mice entering the goal zone in different tasks was measured. All the tasks include (H) task for habituation (T1), and (I) tasks for testing problem-solving skills (T2, T5, T8, and T11), (J) tasks for evaluating short-term (T3, T6, T9, and T12) and (K) long-term memories (T4, T7, T10, and T13). n = 8–11 per group. Data are expressed as mean ± S.E.M. Differences were assessed by two-way ANOVA and Bonferroni’s post hoc test denoted as follows: \**p* < 0.05 and \*\**p* < 0.01 compared with the control group.

The puzzle box test was conducted to further assess the rodent’s problem-solving skill and short-term and long-term memories. Throughout habituation, no notable difference was detected between the laparotomy and control groups, regardless of the mouse strain (Fig. 6h). Between the mouse strains, the PKR^-/-^ mice demonstrated greater latency to enter the goal zone than the wild-type mice (*p* < 0.01; Fig. 6h). In terms of problem-solving ability, the PKR^-/-^ laparotomy mice performed similarly to their PKR^-/-^ control counterparts (Fig. 6i). This finding differed from that observed in the wild-type laparotomy mice who took longer to solve task T2, T5, and T11 (*p* < 0.05; Fig. 6i). In terms of native problem-solving ability, significant differences were also detected between the PKR^-/-^ and wild-type mice (*p* < 0.01; Fig. 6i). In T2, the PKR^-/-^ laparotomy mice demonstrated greater latency when compared to the wild-type laparotomy group (*p* < 0.05; Fig. 6i); a similar trend was also detected in T11 between the PKR^-/-^ controls and their wild-type control counterpart (*p* < 0.05; Fig. 6i). Furthermore, laparotomy did not impair the short-and long-term memories of the PKR^-/-^ mice, and the wild-type laparotomy group showed significant impairment (T7, T12, and T13: *p* < 0.05; T10: *p* < 0.01; Fig. 6j, k). This result suggested that genetic deletion of PKR could ameliorate the cognitive deficits induced by laparotomy.

### Inhibition of PKR in cholinergic neurons ameliorated laparotomy-induced cognitive dysfunctions

rAAV-DIO-PKR-K296R was intracerebroventricularly injected into the right lateral ventricle of the ChAT-IRES-Cre-eGFP mice to inhibit activation of PKR in cholinergic neurons. rAAV expresses a dominant negative form of PKR, that is, PKR-K296R, under the regulation of a double-floxed inverse orientation (DIO) switch (Fig. S4a). At 4 weeks after the stereotaxic injection of rAAV-DIO-PKR-K296R, the fluorescence intensity of p-PKR (T446) signal and the fluorescence ratio of p-PKR to the eGFP signal were reduced in the rAAV-DIO-PKR-K296R-treated group when compared to the saline-treated control (*p* < 0.05; Fig. S4b, c). Therefore, in ChAT-IRES-Cre-eGFP mice, the expression of PKR-K296R effectively suppressed the activation (phosphorylation) of PKR.

At 4 weeks following stereotaxic injection, mice were randomized into four groups: saline-treated control group given only sevoflurane (SAL-CON), saline-treated group subjected to laparotomy (SAL-LAP), AAV-treated group given only sevoflurane (AAV-CON), and AAV-treated group subjected to laparotomy (AAV-LAP). Within 3 days after laparotomy, the SAL-LAP and AAV-LAP groups lost significant weight (*p* < 0.05; Fig. 7a), similar to the wild-type and PKR^-/-^ mice. On POD 6, their body weight returned to normal, and no remarkable difference was detected across all the groups (Fig. 7a).

**Fig. 7.**
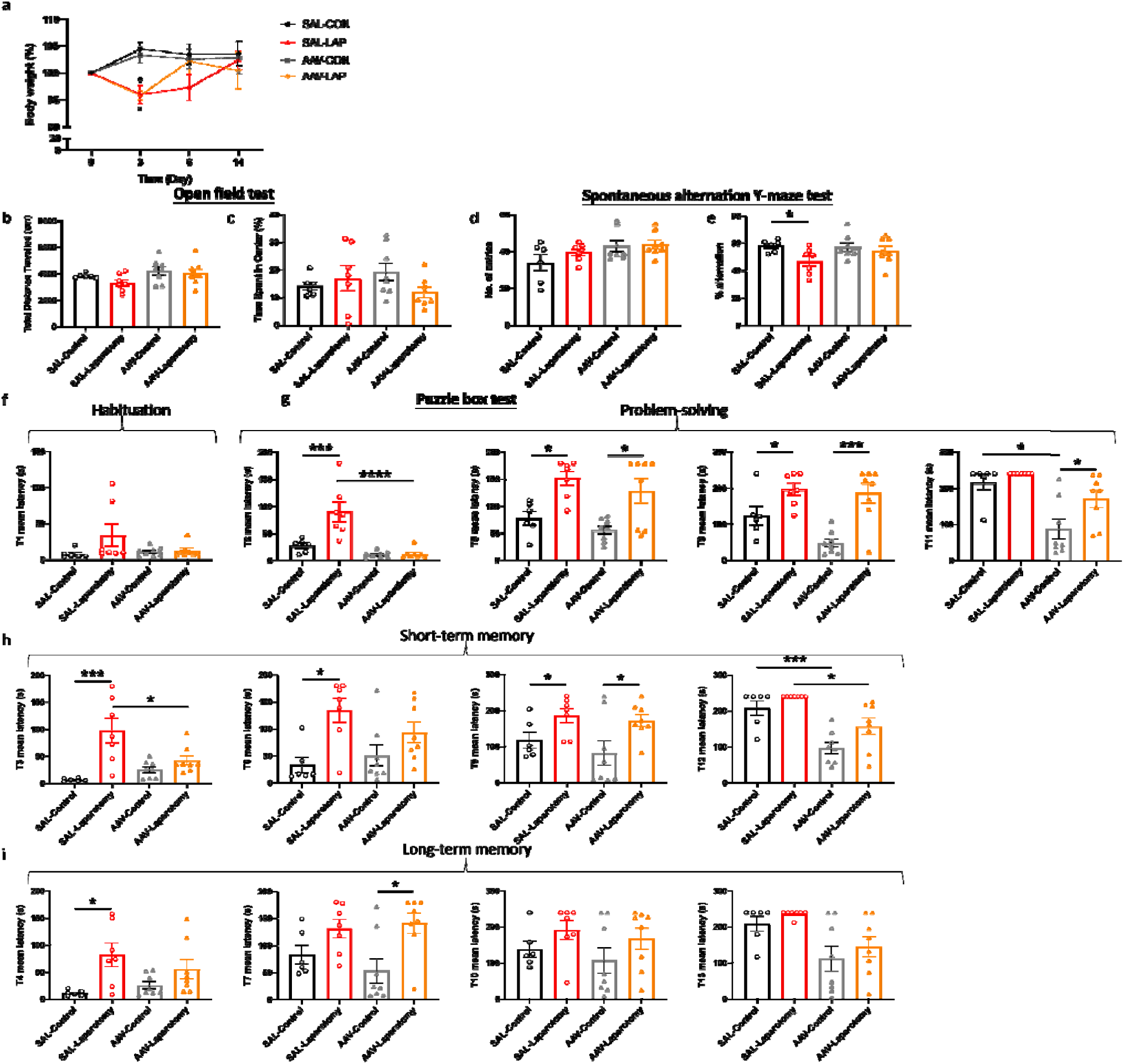
Inhibition of PKR in cholinergic neurons ameliorated laparotomy-induced cognitive impairment. (A) Percentage of the body weight change from baseline of the ChAT-IRES-Cre-eGFP mice of the control group injected with saline (SAL) and the treatment group injected with the rAAV-DIO-PKR-K296R (AAV) throughout the postoperative period. (B) Total distance travelled and (C) central duration of mice in the open-field test. (D) Total number of arm entries and (E) percentage of alternation were measured in the spontaneous alternation Y-maze test. In the puzzle box test, the latency time for the mice entering the goal zone in different tasks was measured. All the tasks include (F) task for habituation (T1), (G) tasks for testing problem-solving skills (T2, T5, T8, and T11), (H) tasks for evaluating short-term (T3, T6, T9, and T12) and (I) long-term memories (T4, T7, T10, and T13). n = 6–8 per group. Data are expressed as mean ± S.E.M. Differences were assessed by two-way ANOVA and Bonferroni’s post hoc test denoted as follows: \**p* < 0.05, \*\**p* < 0.01, \*\*\**p* < 0.001, \*\*\*\**p* < 0.0001 compared with the control group.

Different behavioral tests were conducted to investigate the effects of inhibiting PKR in cholinergic neurons on cognition following laparotomy. rAAV-DIO-PKR-K296R injection and laparotomy did not affect locomotor activity (total distance travelled) and anxiety level (central duration) (Fig. 7b, c). In the SYM test, impaired spatial memory with lower percentage of alternation was detected in the SAL-LAP group when compared to the SAL-CON (*p* < 0.05; Fig. 7e); AAV-treated mice showed no such difference. No remarkable difference in the total number of entries was detected among all groups (Fig. 7d). Our findings suggested that inhibiting PKR in cholinergic neurons could alleviate the spatial working memory deficits induced by laparotomy.

Considering the habituation task (T1) of the puzzle box test, all the groups of mice completed the escape task in about the same amount of time (Fig. 7f). In terms of problem-solving ability, contrasting behaviors were observed. In the T2 task, the SAL-LAP group demonstrated increased latency when compared to the SAL-CON (*p* < 0.001; Fig. 7g); no notable difference was found in the AAV-treated group (Fig. 7g). Meanwhile, the AAV-LAP group solved the T2 task faster than the SAL-LAP group (*p* < 0.0001; Fig. 7g). This result revealed that inhibiting PKR in cholinergic neurons could improve the cognitive functions of mice in the T2 task following laparotomy. In the T5 and T8 task, however, the AAV-treated and saline-treated mice showed increased latency time following laparotomy when compared to their control counterparts (*p* < 0.05; Fig. 7g). With respect to T11 task, the AAV-LAP group demonstrated a significant prolonged latency when compared to the AAV-CON (*p* < 0.05; Fig. 7g). Meanwhile, the AAV-CON group demonstrated better problem-solving skills than the SAL-CON (*p* < 0.05; Fig. 7g).

Regarding short-term memory, consistent results were observed in the T3 and T6 tasks (Fig. 7h). When compared with the SAL-CON group, the SAL-LAP mice showed impaired short-term memory with increased latency to solve the T3 (*p* < 0.001; Fig. 7h) and T6 tasks (*p* < 0.05; Fig. 7h). Such differences were not detected in the AAV-treated groups. In the T3 task, the AAV-LAP group escaped faster than the SAL-LAP group (*p* < 0.05; Fig. 7h). This finding indicated that inhibiting PKR in cholinergic neurons helped alleviate the short-term memory deficits induced by laparotomy. However, for the T9 task, the AAV-treated and saline-treated mice showed increased latency following laparotomy when compared to the controls (*p* < 0.05; Fig. 7h). Moreover, the AAV-treated group demonstrated reduced latency for the T12 task compared with their saline-treated counterpart (*p* < 0.05; Fig. 7h). Therefore, inhibition of PKR in cholinergic neurons could improve short-term memory.

For long-term memory, contrasting behaviors were observed in the T4 and T7 tasks. In the T4 task, the SAL-LAP group showed impairment (*p* < 0.05; Fig. 7i); AAV-treated mice showed no such difference (Fig. 7i). In T7 task, a noticeable change was observed only in the AAV-treated groups (*p* < 0.05; Fig. 7i) but not in the saline-treated groups (Fig. 7i). No significant difference for all the tasks testing long-term memory (T4, T7, T10, and T13 tasks) was observed between the AAV-treated and SAL-treated groups. Therefore, no concrete conclusions concerning the influence of PKR inhibition in cholinergic neurons on long-term memory could be drawn.

### Inhibition of PKR in cholinergic neurons alters glucose metabolism in the frontal cortex in a laparotomy model

MS-based targeted metabolomics was performed to examine the effects of PKR inhibition in cholinergic neurons on brain glucose metabolism. Distinct clustering was observed between the SAL-LAP and AAV-CON groups (Fig. 8a, b). The heatmap depicted the changes of quantified metabolites including tricarboxylic acids and glycolysis intermediates in the frontal cortex across all the groups (Fig. 8c).

**Fig. 8.**
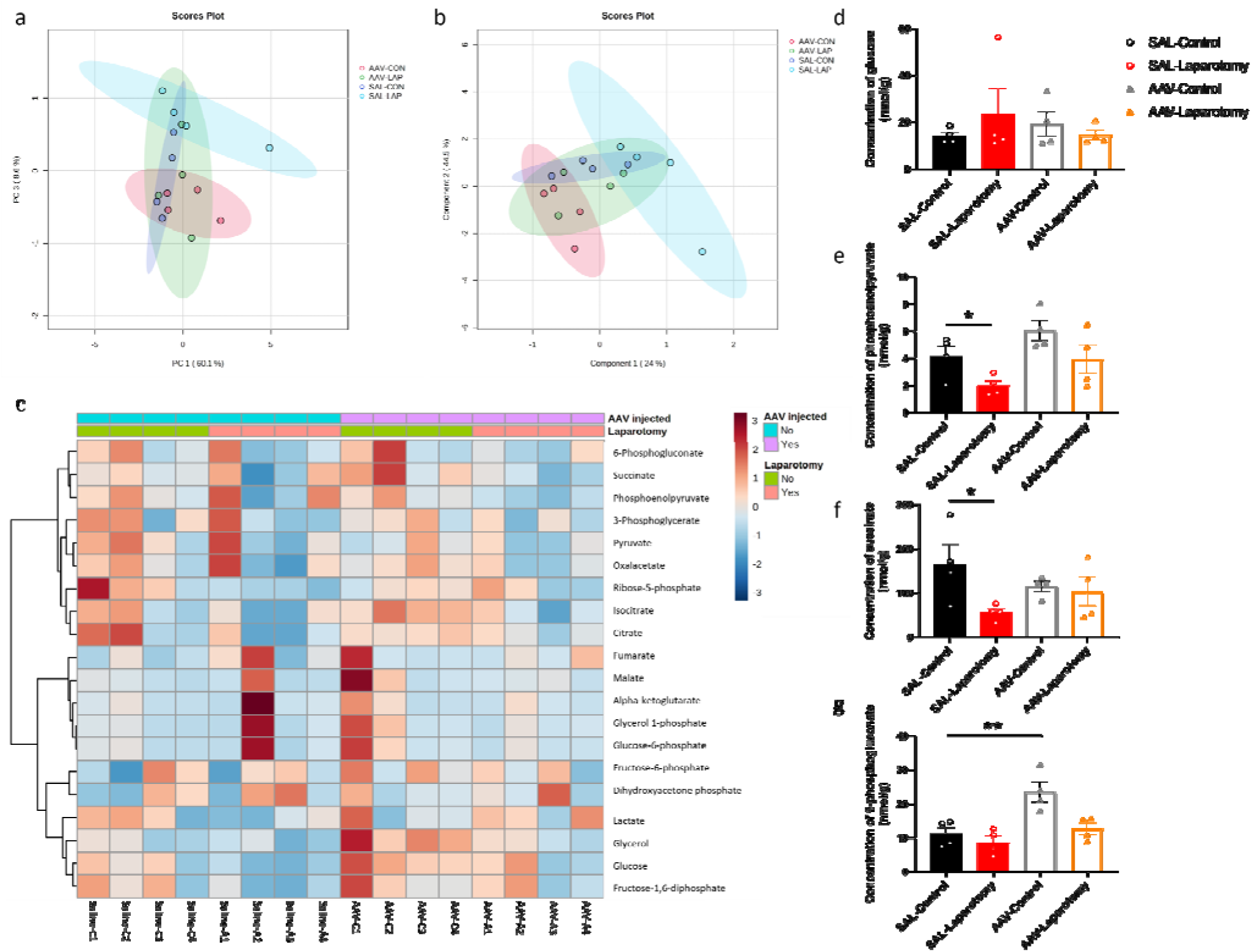
Inhibition of PKR in cholinergic neurons ameliorated laparotomy-induced changes in glucose metabolism in the frontal cortex. Two-dimensional score plots using the (A) principal component analysis (PCA) and (B) partial least squares discriminant analysis (PLS-DA). (C) Heatmap of the metabolites involved in central carbon metabolism in the frontal cortex of the ChAT-IRES-Cre-eGFP mice injected with saline (SAL) or rAAV-DIO-PKR-K296R (AAV). Both groups of mice were then randomly assigned into the laparotomy and control groups. The effects of AAV treatment and laparotomy on glucose metabolism were analyzed using gas chromatography-tandem mass spectrometry (GC-MS/MS). The column represents the samples, and the row displays the glucose metabolites. The heatmap scale ranges from −3 to 3. The brightness of each color corresponded to the magnitude of the log2 fold change of the measured metabolites. Concentration of (D) glucose, (E) phosphoenolpyruvate, (F) succinate, and (G) 6-phosphogluconate were quantified in nmol/g. n = 4 per group. Data are expressed as mean ± S.E.M. Differences were assessed by two-way ANOVA and Bonferroni’s post hoc test denoted as follows: \**p* < 0.05, \*\**p* < 0.01, compared with the control group.

Laparotomy significantly reduced the level of phosphoenolpyruvate in the SAL-LAP group when compared to the control counterpart (*p* < 0.05; Fig. 8e). However, no notable difference in phosphoenolpyruvate level was detected in the AAV-treated group following laparotomy. Therefore, inhibiting PKR in cholinergic neurons could ameliorate the laparotomy-induced changes in phosphoenolpyruvate level. Similarly, laparotomy reduced the succinate level in the SAL-treated mice (*p* < 0.05; Fig. 8f) but not in the AAV-treated group (Fig. 8f). No notable difference in glucose levels was detected among all the groups (Fig. 8d). Apart from the laparotomy-induced changes in glycolysis intermediates, inhibition of PKR in cholinergic neurons significantly elevated the level of 6-phosphogluconate in the AAV-CON group when compared to that in the SAL-CON group (*p* < 0.01; Fig. 8g), suggesting that PKR inhibition in cholinergic neurons could alter brain energy metabolism.

## Discussion

Neuroinflammation is a major cause of cognitive impairment in PND. In this study, we employed a clinically relevant mouse model of laparotomy to induce peripheral and neural inflammation and further demonstrate the regulatory role of PKR in systemic inflammation-induced neuroinflammation and cognitive dysfunctions.

Regarding the laparotomy model, systemic and neural immune responses developed rapidly after the surgery. In theory, laparotomy results in peritoneal air exposure [33, 34], tissue injury, and gut mucosa barrier damage, resulting in increased inflammatory mediators and gut bacteria/endotoxin translocation to the circulation, contributing to systemic inflammation [35]. Our study found a significant elevation in the cytokine mRNA expression of liver (IL-1β and MCP-1) and brain tissues (IL-1β) at four hours after laparotomy (Fig. 1a). On POD 1, laparotomy triggered a significant upregulation of IL-1β and IL-6 expression in frontal cortex and hippocampus (Fig. 1b). IL-1β plays an important role in neuroinflammation and pathogenesis of AD [36] and PD [37]. Through activating nuclear factor κB (NF-κB), IL-1β promotes the upregulation of proinflammatory cytokines including IL-6 and TNF-α [38]. This finding may partially explain the increased IL-6 expression in the brain of the laparotomy group on POD 1 (Fig. 1b). However, the induced systemic inflammation is relatively transient, with proinflammatory cytokine expression levels in the liver returning to normal on POD 1 (Fig. 1b). Therefore, a transient systemic inflammation is already sufficient to induce neuroinflammation as evidenced by the persistent microgliosis revealed by flow cytometry (Figs. 1c–f) and confocal imaging (Fig. 2) on POD 14. Our findings are also consistent with a prior study that demonstrated significant microglial activation in frontal cortex and hippocampus following laparotomy [12]. Beyond the microglial activation, cognitive deficits (Fig. 3) were also detected in the wild-type laparotomy model mice. Therefore, the laparotomy model is a viable and relevant paradigm for studying systemic inflammation-triggered neuroimmune response and cognitive dysfunction.

Our study is the first to show that genetic deletion of PKR could alleviate laparotomy-induced peripheral and neural inflammation, as well as downregulate the increased expression of IL-1β and IL-6 in liver, frontal cortex, and hippocampus following laparotomy (Fig. 4a and b). Since PKR plays a critical role in IL-1β activation, PKR knockout can help prevent pro-IL-1β from cleaving into IL-1β. These findings were consistent with other previous studies showing that PKR could attenuate inflammation by regulating inflammasome activation [39] and various inflammatory pathways including TNF [40], p38 MAPK [21], and NF-κB/IκB signaling pathway [41]. PKR inhibition prevented IκB and NF-κB activation and decreased IL-1β expression in primary murine mixed neuron–astrocyte–microglia co-culture model [41] and RAW264.7 macrophages [42]. Subsequently, the reduced IL-1β expression further downregulated IL-6 expression by preventing the activation of NF-κB. Four hours following surgery, a significant elevation was found in TNF-α expression in the liver of the PKR^-/-^ laparotomy group but not in their wild-type counterpart (Fig. 4a). However, no notable difference was detected between the PKR^-/-^ and wild-type laparotomy groups. Similar findings were also observed in our earlier investigation, which demonstrated a significant elevation in TNF-α expression in the hippocampus of the PKR^-/-^ mice after *Escherichia coli* challenge but not in their wild-type control counterpart [16]. Indeed, PKR is involved in TNF-dependent activation of c-Jun N-terminal kinase (JNK), NF-kB and protein kinase B. Meanwhile, knockout of PKR could abrogate the TNF-induced activation of the above signaling pathways. Therefore, the PKR^-/-^ laparotomy mice may tend to highly express TNF-α to activate the downstream pathways. Our study has taken a significant step beyond previous research. Although previous works showed that genetic deletion of PKR in mice can prevent activation of microglia following intraperitoneal injection of LPS [43], we demonstrated that PKR also alleviates neuroinflammation induced by sterile systemic inflammation as evidenced by the suppressed laparotomy-induced microglial activation in frontal cortex and hippocampus of PKR^-/-^ mice (Fig. 4, 5). Therefore, PKR regulates the activation of microglia/macrophages. Since PKR is critical for LPS-induced inducible nitric oxide synthase expression [44], it is responsible for the macrophage inflammatory response by regulating the production of a proinflammatory factor, nitric oxide [45]. Moreover, PKR is necessary for the dsRNA-induced activation of macrophages [42]. All the aforementioned studies supported the role of PKR in activation of microglia/macrophages. Furthermore, knockout of PKR in mice could rescue the laparotomy-triggered impairment of problem-solving ability, spatial working memory, and short-term and long-term memories in mice (Figs. 6e, g, and i–k).

Another novel finding of this project is that inhibition of PKR in cholinergic neurons could rescue the laparotomy-induced cognitive deficits. The cholinergic system is selected for investigation because it is not only essential for cognition but also regulates inflammation through the cholinergic anti-inflammatory pathway [46], which connects systemic inflammation and neuroinflammation. This would elucidate the neuron-specific effects of PKR on cognition during inflammation beyond its immune regulatory roles. This study also demonstrated that inhibition of PKR in cholinergic neurons could alleviate the laparotomy-induced impairment of short-term and spatial working memory (Fig. 7e and h). A previous study showed that PKR inhibition could improve cortex-dependent memory consolidation in rats and mice [47]. Meanwhile, PKR^−/−^ mice demonstrated enhanced conditioned taste aversion memory when compared with their wild-type control [47]; while PKR activation could impair hippocampal late-phase LTP and memory consolidation in PKR recombinant mice [48]. All the abovementioned studies supported the crucial role of PKR in regulating cognitive functions. Our research highlights PKR as a potential key target in treating inflammation-induced cognitive deficits, opening up promising possibilities for future therapeutic approaches focusing on PKR modulation.

We observed that laparotomy could lead to brain glucose hypometabolism with reduction in glycolysis and citric acid cycle intermediates (phosphoenolpyruvate and succinate) (Fig. 8), which is a hallmark of neurodegenerative diseases including AD and PD in preclinical stages [49, 50]. Progressive reduction in brain glucose metabolism positively correlates with amyloid load and cognitive decline in patients with early AD or PD suffering from mild cognitive impairment [50–52]. AD patients showed reduced concentrations of glycolytic intermediates including phosphoenolpyruvate in cerebrospinal fluid [53]. By contrast, administering succinate into the brain by retrodialysis could improve the brain reduced/oxidized nicotinamide adenine dinucleotide redox state [54] and brain energy metabolism in patients suffering from traumatic brain injury [55]. All of the abovementioned studies established a close association between brain glucose metabolism and cognition.

Our study unraveled the novel role of PKR in regulating brain glucose metabolism in the laparotomy model. Blocking PKR in cholinergic neurons could rescue the laparotomy-induced alterations in glucose metabolism in the frontal cortex on POD 14 (Fig. 8e and f). PKR regulates glucose homeostasis [56] and induces insulin resistance [57, 58]. Increased eIF2α phosphorylation is a hallmark of insulin resistance and obesity [57–59]. Under metabolic and obesity-induced stress, PKR could form a complex with TAR RNA-binding protein (TRBP), activating JNK [56]. Inhibiting TRBP in the liver could improve insulin sensitivity, glucose metabolism, and alleviate inflammation [56]. The interaction between PKR and TRBP links metabolism to inflammatory signaling regulation. However, no prior study has examined the effect of blocking PKR in brain on glucose metabolism. Our findings supported that PKR in cholinergic neurons play a critical role in modulating glucose metabolism. Further study to unveil the underlying regulatory networks is warranted.

## Conclusions

The laparotomy model was a stable and viable model for studying neuroimmune responses triggered by systemic inflammation as evidenced by microgliosis and cognitive dysfunction. By genetic deletion of PKR, we demonstrated the critical role of PKR in regulating peripheral and neural inflammation. Knockout of PKR could help alleviate neuroimmune responses and cognitive impairment induced by laparotomy. Furthermore, inhibition of PKR in the cholinergic neurons of the laparotomy mice demonstrated a neuroprotective role and improved the glucose metabolism and cognitive functions. On the basis of our combined findings, PKR could be a pharmacological target for treating systemic inflammation-induced neuroinflammation and cognitive deficits.

## List of abbreviations

AAV-CON: The AAV-treated group given only sevoflurane
AAV-LAP: The AAV-treated group subjected to laparotomy
AD: Alzheimer’s disease
CA: Cornu ammonis
CD68-eGFP mice: C57BL/6-Tg(CD68-EGFP)1Drg/J mice
DAPI: 4’, 6-Diamidino-2-phenylindole
DG: Dentate gyrus
DIO: Double-floxed inverse orientation
dsRNA: Double-stranded RNA
eGFP: Enhanced green fluorescent protein
eIF2α: Eukaryotic initiation factor-2alpha
GC-MS/MS: Gas chromatography-tandem mass spectrometry
Iba1: Ionized calcium-binding adaptor molecule 1
JNK: c-Jun N-terminal kinase
LPS: Lipopolysaccharide
LTP: Long-term potentiation
MAPKs: Mitogen-activated protein kinases
MCP: Monocyte chemoattractant protein
NF-κB: Nuclear factor
κB NOR: Novel object recognition test
OFT: Open field test
PD: Parkinson’s disease
PKR: Double-stranded RNA-dependent protein kinase
PND: Perioperative neurocognitive disorders
POD: Postoperative day
SAL-CON: The saline-treated control group given only sevoflurane
SAL-LAP: The saline-treated group subjected to laparotomy
SYM: Spontaneous alternation Y-maze test
TMEM119: Transmembrane protein 119
TRBP: TAR RNA-binding protein

## Conflict of Interest

Authors declare no competing interests.

## Ethics approval and consent to participate

All experiments were approved by the Department of Health of the Hong Kong SAR government and the CULATR of the University of Hong Kong (#4911-18 and #5819-21).

## Availability of data and materials

The online version contains supplementary material. All raw data are available at the HKU data repository.

## Funding

This study is supported by General Research Fund (17123217).

## Notes

### Competing Interest Statement

The authors have declared no competing interest.

### Summary of Updates

The manuscript has been revised to focus more on the discussion of the role of PKR in modulating glucose metabolism and cognitive functions in cholinergic neurons.

