## Supplementary material for "Regulatory Role of PKR in Systemic Inflammation-Triggered Neuroinflammation and its Modulation of Glucose Metabolism and Cognitive Functions in Cholinergic Neurons": Figure S1-4; Table S1-3

**Supplementary Methods**

*RNA isolation and reverse transcription*

Total RNA was extracted from the tissue samples using RNAiso Plus (TAKARA, Japan) according to the manufacturer’s protocol. In brief, the samples were homogenized with RNAiso Plus and incubated for 5 min before adding chloroform. After 5 min of incubation, samples were centrifuged for 15 min at 12 000 x *g* (4°C). Isopropanol was added to the upper aqueous phase and mixtures were then centrifuged 10 min at 12 000 x *g* (4°C). The precipitated RNA pellets were rinsed with ice-cold 75% ethanol before centrifugation for 5 min at 7500 x *g* (4°C). The pellets were then air-dried and then resuspended in RNase-free water containing 0.1 mM EDTA. The concentration of the RNA samples was measured by the Nanodrop One spectrophotometer (Thermo Scientific, USA). The RNA samples with the ratio of A260/A280 larger than 1.8 and A260/A230 ratio in the range of 2.0-2.2 were proceeded for analysis. RNA samples were then reverse transcribed into cDNA using the PrimeScript™ RT Master Mix (TAKARA, Japan) based on the manufacturer’s protocol. cDNA samples were then stored at -20°C.

*Cryosectioning*

After post-fixation of brain samples for 24 h at 4% paraformaldehyde (PFA), the tissues were dehydrated overnight at 4°C in a 3-step sucrose gradient (20%, 30% and 30% sucrose in PBS). Samples were then embedded in Tissue-Tek optimal cutting temperature (O.C.T.) medium (Sakura Finetek, USA) and frozen at -20ºC prior to cryosection. The frozen sample blocks were then sectioned on a cryostat (Leica) at 20 µm for immunostaining. Sample slides were then air-dried and stored at -80ºC.

*Sample Preparation for targeted metabolomics*

The frontal cortex samples were collected and weighed at the end of the experiment. After homogenizing 50 mg of tissue sample in 500 μl of methanol/water (80%, v/v), 200 ng norvaline was added as an internal standard, with the addition of 250 µl of 0.1 M HCl. The sample mixture was than vortexed for 30 s. To extract the metabolites, sample was then mixed with 400 µl of chloroform. The sample was agitated for 15 min before being centrifuged for 5 min at 16,000 x *g* at 4°C. Before derivatization, 375 μl of supernatant was then dried at room temperature under nitrogen. A two-step derivatization procedure was applied. In the first step, the dried residue was redissolved and derivatized with 40 μl of methoxylamine hydrochloride (30 mg/ml in pyridine) for 2 h at 37°C to convert keto- and aldehyde groups. In the second step, the sample was trimethylsilylated in 70 μl N-Methyl-N-(trimethylsilyl)trifluoroacetamide (MSTFA) with 1% trimethylchlorosilane (TMCS) for 1 h at 37°C.

*Targeted metabolomic by GC-MS/MS*

Targeted metabolomics on central carbon metabolism was analyzed using gas chromatography-tandem mass spectrometry (GC-MS/MS). GC-MS/MS analysis was performed on Agilent 7890B GC-Agilent 7010 Triple Quadrupole Mass Spectrometer system (Agilent, USA) using SCAN and multiple reaction monitoring (MRM) mode. One µl of sample was injected for GC-MS/MS analysis. An Agilent DB-5MS capillary column (30 m × 0.25 mm, 0.25 µm film thickness) (Agilent, USA) was used in the GC to separate the sample under constant flow at 1 mL/min. The GC oven was programmed from 60°C (hold time: 1 min) to 120°C at 10°C/min, then to 150°C at 3°C/min, and then to 200°C at 10°C/min and finally to 280°C at 30°C/min (hold time: 5 min). The inlet temperature and transfer line temperature were maintained at 250°C and 280°C respectively. In MRM mode, two characteristic transitions, including quantifier and qualifier transitions, were monitored. Mass spectrometry data were acquired in the SCAN mode (mass range, m/z 50-500). Mass spectrometry data were analyzed using the Agilent MassHunter Workstation quantitative analysis software. For each analyte, the linear calibration curve was generated by plotting peak area ratio of external/internal standard against standard concentration at various concentration levels. The analytes were then validated by comparing the retention time and ratio of characteristic transitions between the sample and standard. Data were then normalized by median and log-transformed. Heat map and hierarchical clustering were performed based on Euclidian distance using Ward’s method. Principal component analysis (PCA) and partial least squares discriminant analysis (PLS-DA) were further performed using Metaboanalyst 5.0.

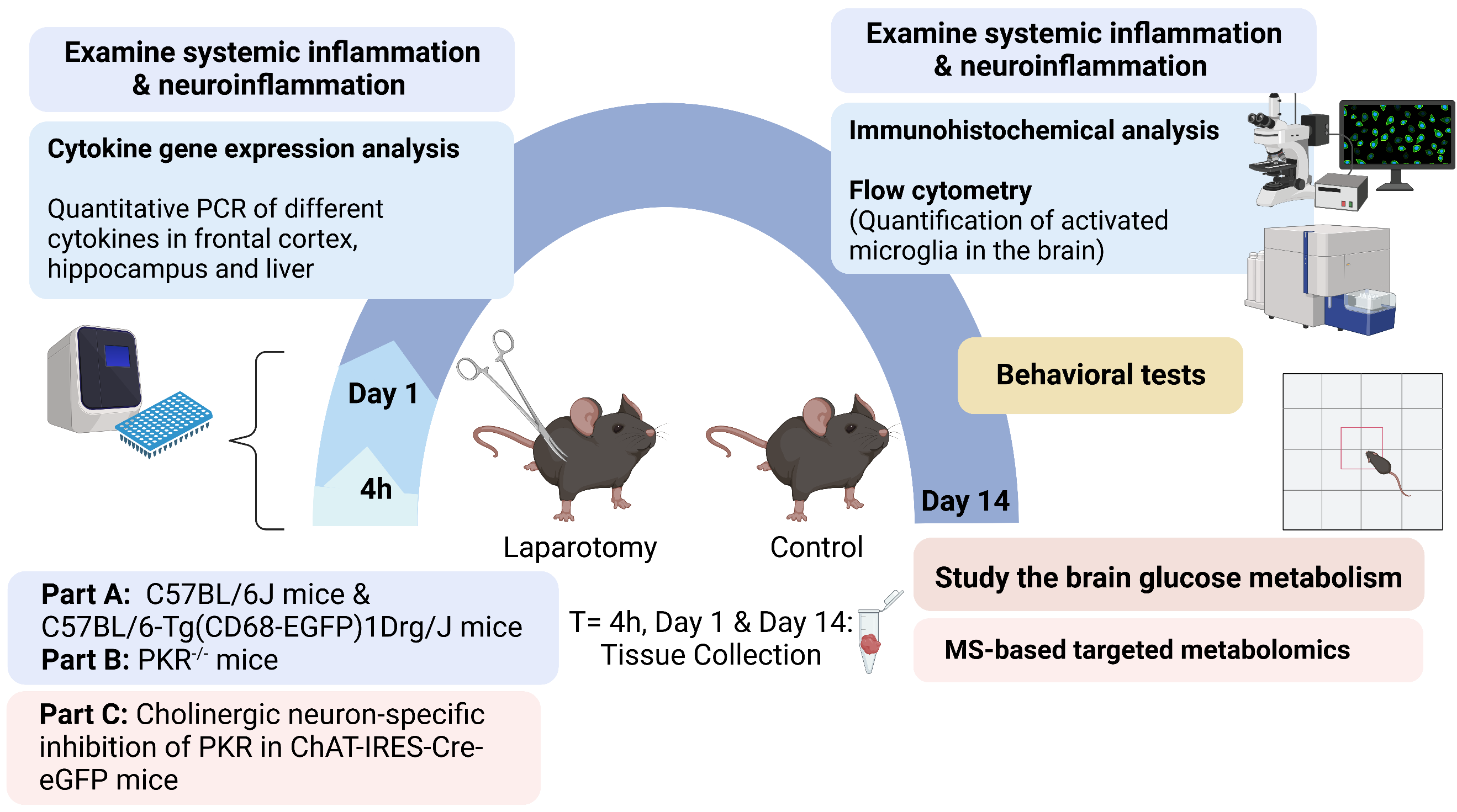

**Figure S1. The experimental workflow.** For the first part, both male wild-type C57BL/6J and C57BL/6-Tg(CD68-EGFP)1Drg/J mice were assigned into 2 groups randomly: laparotomy under sevoflurane anesthesia and control group under sevoflurane anesthesia. In the second part, PKR^-/-^ mice were exposed to laparotomy with sevoflurane anesthesia or sevoflurane anesthesia to examine the role of PKR in regulating systemic inflammation-triggered neuroinflammation. For the third part, intracerebroventricular injection of rAAV-DIO-PKR-K296R into the right lateral ventricle of ChAT-IRES-Cre-eGFP mice was performed to inhibit PKR activation in cholinergic neurons. Mice were then subjected to laparotomy with sevoflurane or sevoflurane anesthesia respectively. The effects of blocking PKR in cholinergic neurons on modulating glucose metabolism and cognitive functions were examined in the laparotomy model.

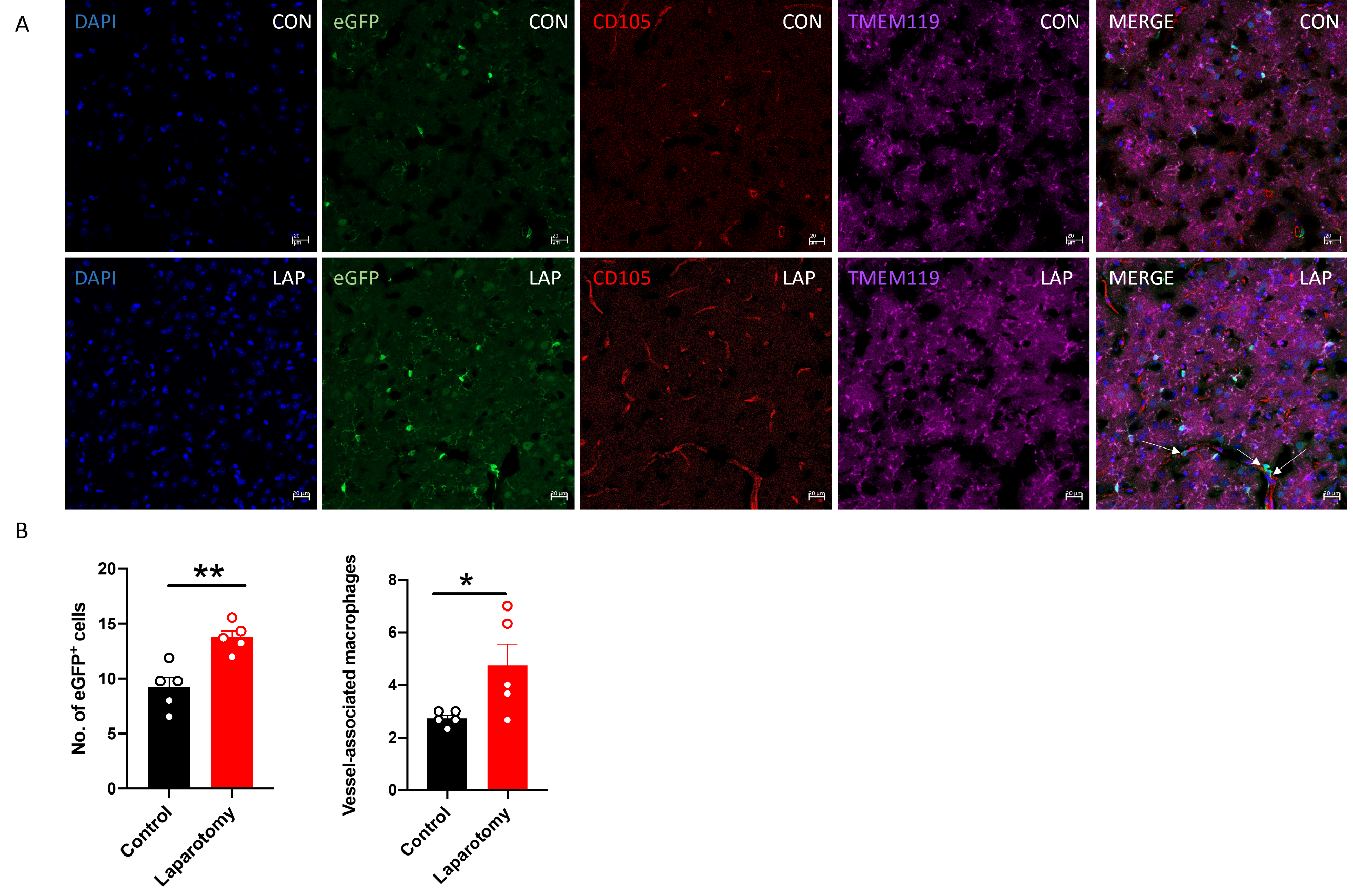

**Figure S****2. Activation of macrophage in the frontal cortex following laparotomy.** (A) Representative confocal images of the immunohistochemical staining of DAPI, eGFP, CD105 and TMEM119 in frontal cortex sections from CD68-eGFP mice of the control and laparotomy group on postoperative day 14. The white arrows represent vessel-associated macrophages. (B) The number of eGFP positive cells and number of vessel-associated macrophages were quantified. n = 5 per group. Data are expressed as mean ± S.E.M. Differences were assessed by unpaired two-tailed Student’s t-test denoted as follows: ^*^*p* < 0.05 and ^**^*p* < 0.01 compared with the control group.

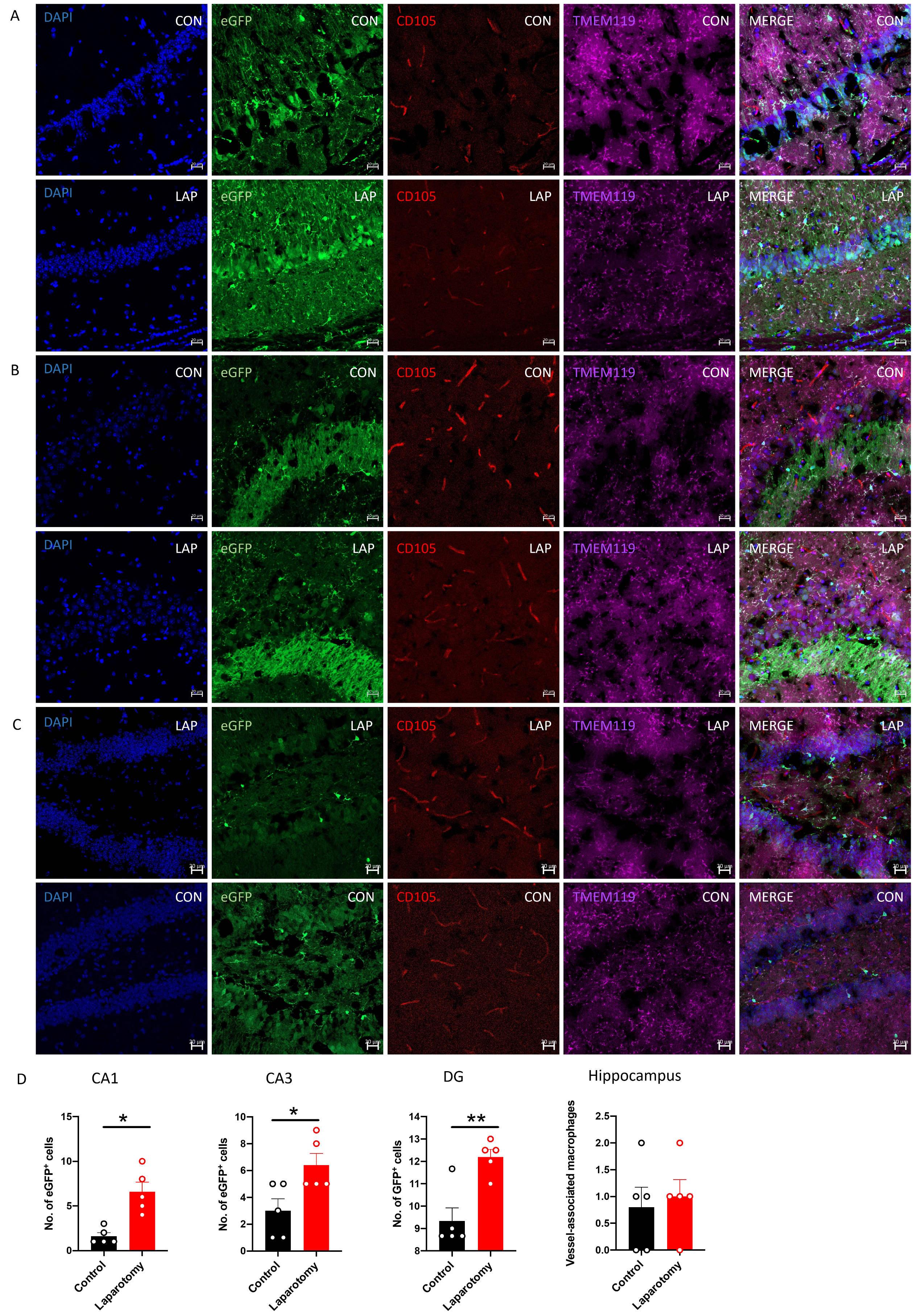

**Figure S3. Activation of macrophage in the hippocampus following laparotomy.** Representative confocal images of the immunohistochemical staining of DAPI, eGFP, CD105 and TMEM119 in the (A) cornu ammonis (CA) 1, (B) CA3 and (C) dentate gyrus (DG) of the hippocampus sections from CD68-eGFP mice of the control and laparotomy group on postoperative day 14. (D) The number of eGFP positive cells and number of vessel-associated macrophages were quantified. n = 5 per group. Data are expressed as mean ± S.E.M. Differences were assessed by unpaired two-tailed Student’s t-test denoted as follows: ^*^*p* < 0.05 and ^**^*p* < 0.01, compared with the control group.

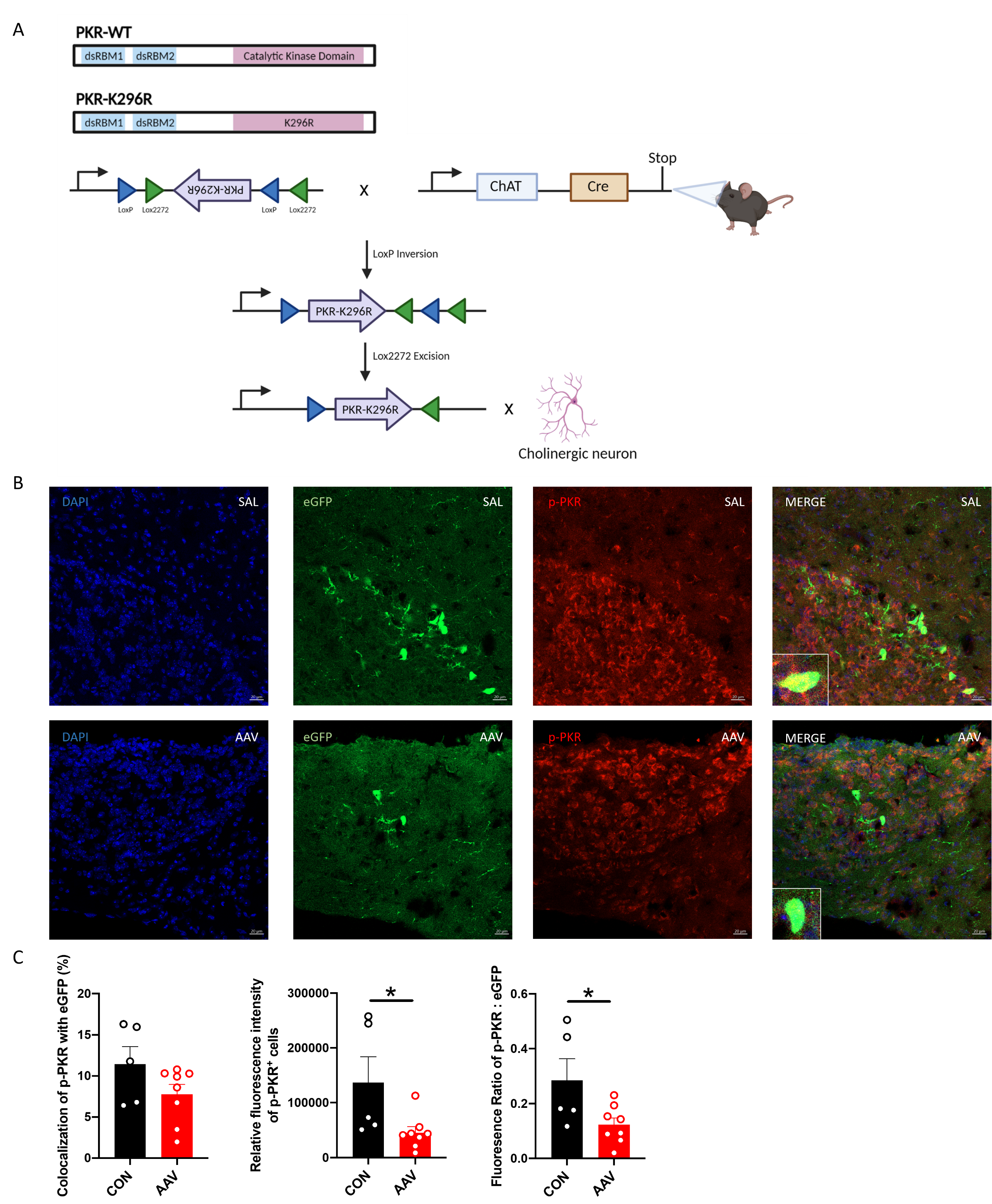

**Figure S4. Stereotaxic injection of rAAV-DIO-PKR-K296R inhibited the phosphorylation of PKR on cholinergic neurons of the ChAT-IRES-Cre-eGFP mice.** (A) Schematic diagrams illustrating the PKR-K296R construct. The PKR-K296R, which is an unphosphorylated kinase-dead PKR molecule, has a catalytically inactive mutation in the ATP binding site. In the absence of Cre, the inverted PKR-K296R are inactivated. Upon the administration of rAAV-DIO-PKR-K296R into the mouse brain with ChAT-IRES-Cre, the PKR-K296R will be activated in the cholinergic-neurons. (B) Representative confocal images of the immunohistochemical staining of DAPI, eGFP and p-PKR in the medial habenula from the ChAT-IRES-Cre-eGFP mice of the control group injected with saline (SAL) and the treatment group injected with rAAV-DIO-PKR-K296R (AAV) at 4 weeks after the stereotaxic injection. (C) The percentage of colocalization of the p-PKR with the eGFP signal, the fluorescence ratio of p-PKR to eGFP in cholinergic neurons, and the relative fluorescence intensity of p-PKR^+^ cells in cholinergic neurons were quantified. n = 5–8 per group. Data are expressed as mean ± S.E.M. Differences were assessed by unpaired two-tailed Student’s t-test*,* denoted as follows: ^*^*p* < 0.05, compared with the control group.

**Table S1. Experimental scheme of the puzzle box test.**

| Trial | Day | Condition | Obstruction | Time limit (s) |
| --- | --- | --- | --- | --- |
| T1 | 1 | 0 | Open door with no obstruction | 180 |
| T2 | 1 | 1 | Open channel within doorway | 180 |
| T3 | 1 | 1 | Open channel within doorway | 180 |
| T4 | 2 | 1 | Open channel within doorway | 180 |
| T5 | 2 | 2 | Channel filled with bedding | 180 |
| T6 | 2 | 2 | Channel filled with bedding | 180 |
| T7 | 3 | 2 | Channel filled with bedding | 180 |
| T8 | 3 | 3 | Tissue plug within doorway | 240 |
| T9 | 3 | 3 | Tissue plug within doorway | 240 |
| T10 | 4 | 3 | Tissue plug within doorway | 240 |
| T11 | 4 | 4 | Foam plug within doorway | 240 |
| T12 | 4 | 4 | Foam plug within doorway | 240 |
| T13 | 5 | 4 | Foam plug within doorway | 240 |

**Table S2. Sequences of primers for real-time PCR.**

| Gene | Primer sequence |
| --- | --- |
| IL-1β | F: 5’-GATGAAGGGCTGCTTCCAAAC-3’ |
|  | R: 5'-TCCACAGCCACAATGAGTGA-3' |
| IL-6 | F: 5'-TTCACAAGTCCGGAGAGGAG-3' |
|  | R: 5'-TCCACGATTTCCCAGAGAAC-3' |
| MCP-1 | F: 5'-TGCTGTCTCAGCCAGATGCAGTTA-3' |
|  | R: 5'-TACAGCTTCTTTGGGACACCTGCT-3' |
| TNF-α | F: 5'-CCCCAGTCTGTATCCTCCT-3' |
|  | R: 5'-ACTGTCCCAGCATCTTGT-3' |
| GAPDH | F: 5′-ATTCAACGGCACAGTCAA-3′  R: 5′-CTCGCTCCTGGAAGATGG-3′ |

Abbreviations: IL-1β, interleukin-1β; IL-6, interleukin-6; MCP-1, monocyte chemoattractant protein-1; TNF-α, tumor necrosis factor alpha; GAPDH, glyceraldehyde-3-phosphate dehydrogenase.

**Table S3. Primary antibodies used in immunohistochemical staining.**

| Antibody | Dilution | Brand | Catalog no. |
| --- | --- | --- | --- |
| Goat anti-GFP | 1:150 | Novusbio | Nb-100-1770 |
| Rat anti-CD105 | 1:200 | Thermo Fisher Scientific | 14-1051-82 |
| Rabbit anti-Iba1 | 1:150 | FUJIFILM Wako | 019-19741 |
| Rabbit anti-TMEM119 | 1:200 | Abcam | ab209064 |
| Rabbit Anti-PKR Phospho-T446 | 1: 100 | Abcam | ab32036 |

Abbreviations: GFP, Green fluorescent protein; CD105, endoglin; Iba1, ionized calcium-binding adaptor molecule 1; TMEM119, Transmembrane protein 119; PKR, protein kinase R
